# Neuronal potassium channel activity triggers initiation of mRNA translation through binding of translation regulators

**DOI:** 10.1101/2024.02.07.579306

**Authors:** Taylor J. Malone, Jing Wu, Yalan Zhang, Pawel Licznerski, Rongmin Chen, Sheikh Nahiyan, Maysam Pedram, Elizabeth A. Jonas, Leonard K. Kaczmarek

## Abstract

Neuronal activity stimulates mRNA translation crucial for learning and development. While FMRP (Fragile X Mental Retardation Protein) and CYFIP1 (Cytoplasmic FMR1 Interacting Protein 1) regulate translation, the mechanism linking translation to neuronal activity is not understood. We now find that translation is stimulated when FMRP and CYFIP1 translocate to the potassium channel Slack (KCNT1, Slo2.2). When Slack is activated, both factors are released from eIF4E (Eukaryotic Initiation Factor 4E), where they normally inhibit translation initiation. A constitutively active Slack mutation and pharmacological stimulation of the wild-type channel both increase binding of FMRP and CYFIP1 to the channel, enhancing the translation of a reporter for β-actin mRNA in cell lines and the synthesis of β-actin in neuronal dendrites. Slack activity-dependent translation is abolished when both FMRP and CYFIP1 expression are suppressed. The effects of Slack mutations on activity-dependent translation may explain the severe intellectual disability produced by these mutations in humans.

**HIGHLIGHTS:**

- Activation of Slack channels triggers translocation of the FMRP/CYFIP1 complex
- Slack channel activation regulates translation initiation of a β-actin reporter construct
- A Slack gain-of-function mutation increases translation of β-actin reporter construct and endogenous cortical β-actin
- FMRP and CYFIP1 are required for Slack activity-dependent translation

**IN BRIEF:** Malone *et al*. show that the activation of Slack channels triggers translocation of the FMRP/CYFIP1 complex from the translation initiation factor eIF4E to the channel. This translocation releases eIF4E and stimulates mRNA translation of a reporter for β-actin and cortical β-actin mRNA, elucidating the mechanism that connects neuronal activity with translational regulation.

## INTRODUCTION

Slack (Sequence Like a Calcium-Activated Potassium Channel, also termed K_Na_1.1, KCNT1, or Slo2.2) is a sodium-activated potassium channel (K_Na_ channel) that is predominantly expressed in neurons of the central nervous system where it regulates neuronal excitability^1-5^. In the last ten years, gain-of-function mutations in the Slack channel have been identified in patients with several distinct early onset epilepsy disorders^6-8^, all of which are also associated with severe intellectual disability^9^. Slack was also the first of several ion channels shown to interact directly with the mRNA-binding protein Fragile X Mental Retardation Protein (FMRP)^10,11^. Other FMRP-binding channels include Kv1.2, Kv4.3, BK, SK2, Cav2.2 and Cav3.1^12-18^. In all cases, the interaction of FMRP alters some aspects of channel activity. For example, FMRP binding to Slack increases the Slack open probability and current and modulates action potential shape^10,11^.

Loss of expression of FMRP in humans results in Fragile X Syndrome, the leading inherited cause of intellectual disability and autistic spectrum behavior, with co-occurring seizures, anxiety, hyperactivity, hypersensitivity, and hyperarousal^19,20^. In neurons, FMRP regulates both the trafficking and activity of its targets, controlling local translation of mRNAs, such as those for Map1b, Arc (Activity-Regulated Cytoskeleton-Associated Protein), and CaMKIIα, via several different mechanisms^20-23^. Among its functions, FMRP controls initiation of translation by forming a complex with CYFIP1 (Cytoplasmic FMR1 Interacting Protein 1, also known as an eIF4E-binding protein, 4E-BP)^24^. The FMRP/CYFIP1 complex binds the initiation factor eIF4E (Eukaryotic Initiation Factor 4E), suppressing the first step in protein synthesis. A previous study demonstrated that upon brief neuronal stimulation, the FMRP/CYFIP1 complex dissociates from eIF4E, releasing FMRP target mRNAs for translation^24^.

The regulation of ion channel activity, including that of Slack, by FMRP suggests that Slack could reciprocally regulate the functions of FMRP in translation. This is supported by the finding that the cytoplasmic C-terminal domain of Slack channels also binds FMRP’s partner CYFIP1^25^, and that Slack channels have been found to be required for protein synthesis-dependent changes in the excitability of neurons in *Aplysia*^11^. We have now found that a disease-causing mutation in the C-terminus of Slack (*Slack-R455H*) that constitutively activates the channel, as well as pharmacological stimulation of the wild-type channel, triggers the translocation of FMRP/CYFIP1 from eIF4E to the channel complex. Furthermore, we demonstrate that activation of Slack channels stimulates translation both in cell lines and in neurons, and that the disease-causing *Slack-R455H* mutation results in constitutive activation of translation, an effect that requires the untranslated region (UTR) of the mRNA. Finally, the suppression of FMRP and CYFIP1 expression abolishes the ability of the channel to stimulate translation. The link between channel activity and stimulation of protein synthesis, and its dysregulation by mutations, is likely to play a key role in the intellectual disability observed in patients with Slack mutations.

## RESULTS

### Activation of Slack channels triggers translocation of the FMRP/CYFIP1 complex

Slack channels can be coimmunoprecipitated with FMRP and CYFIP1^10,25^. To determine whether these interactions are altered by Slack mutations that result in severe intellectual disability and early onset seizures, we tested the interaction of both proteins with *Slack-R455H* channels. The *Slack-R455H* mutation in mice corresponds to the human mutation *Slack-R474H*, which causes EIMFS (Epilepsy of Infancy with Migrating Focal Seizures) with severe intellectual disability and results in one the greatest increases in K^+^ current among Slack mutations^26,27^. As is the case with humans, heterozygote mice bearing one copy of this mutation have seizures and persistent interictal spikes but homozygous *Slack-R455H* mice do not survive birth^27^. For simplicity, the heterozygotes are termed *Slack-R455H* mice in this study.

Immunoprecipitation (IP) from extracts of cerebral cortex of 2-month-old wild-type and *Slack-R455H* mutant mice was carried out using an antibody against the N-terminal domain of the Slack channel^28,29^. We found that protein levels of FMRP and CYFIP1 immunoprecipitated by Slack were higher in cortical lysates from *Slack-R455H* mice than wild-type mice, while there were no differences in the levels of total or immunoprecipitated Slack in the two conditions (**Fig. 1A,C**). Higher levels of phosphorylated FMRP (p-FMRP, a post-translation modification suggested to regulate its activity^30,31^) were also associated with *Slack-R455H* mutant channel. To further determine if the increased binding of FMRP/CYFIP1 to the Slack channel altered the amounts of FMRP/CYFIP1 bound to the translation initiation factor eIF4E, IP was carried out using an anti-eIF4E antibody. We found that the levels of FMRP, p-FMRP and CYFIP1 immunoprecipitated by eIF4E was significantly decreased in cortical lysates from *Slack-R455H* mice compared with those from wild-type mice (**Fig.1B,C**). There were no differences in levels of eIF4E between the two conditions.

**Figure 1:**
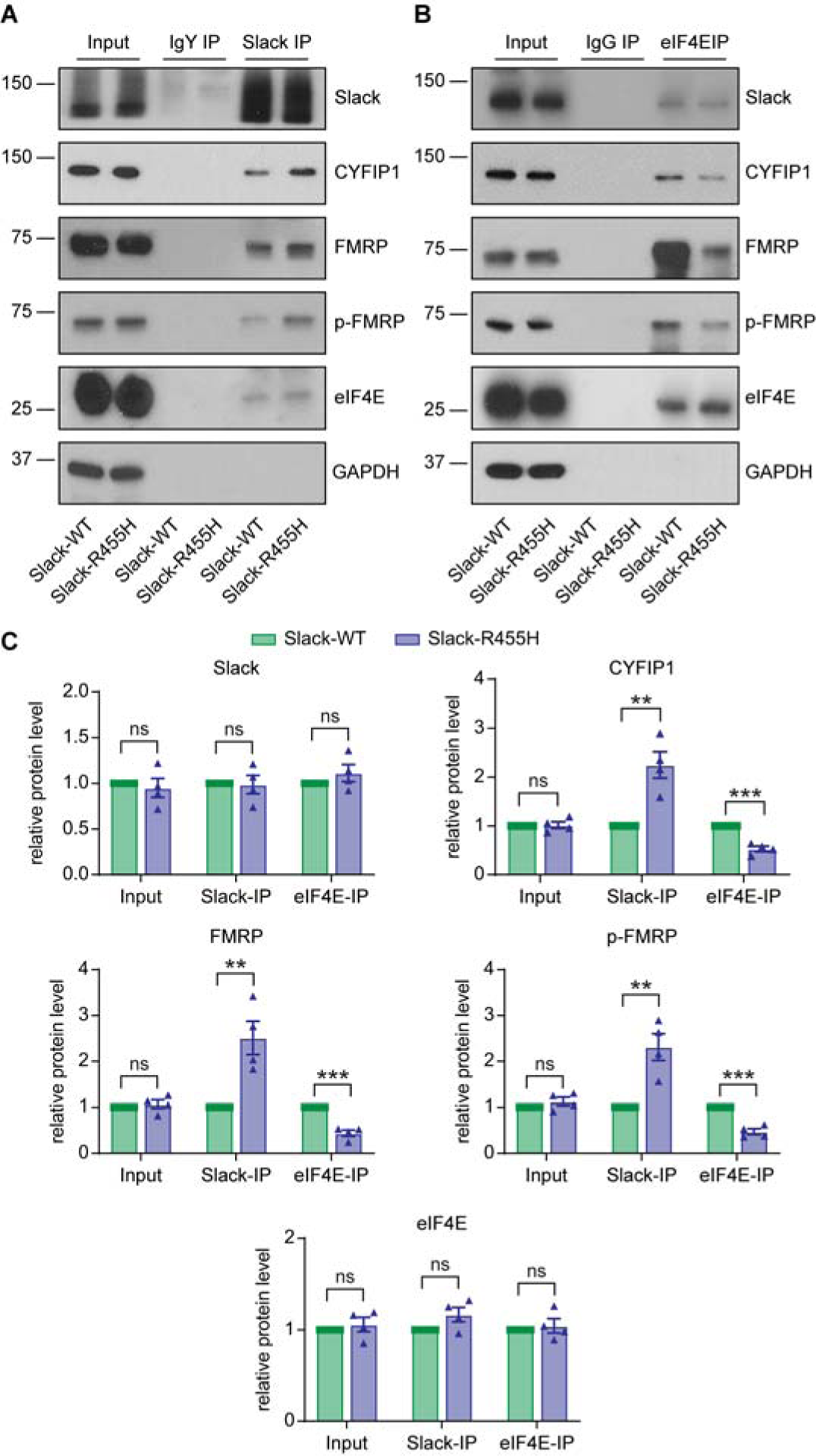
*Slack-R455H* mutation triggers translocation of the FMRP/CYFIP1 complex. (A, B) Immunoprecipitation of Slack (A) or eIF4E (B) from mouse cerebral cortex. (C) Quantification of western blots in a and B. Input, Slack IP, and eIF4E IP samples are normalized by GAPDH, Slack, and eIF4E levels, respectively, followed by normalization to *Slack-WT* level (n=4 mice, paired t-test). Data are expressed as mean ± SEM. *\*p*<0.05, *\*\*p*<0.005, *\*\*\*p*<0.0005

We next tested whether translocation of FMRP/CYFIP1 to the Slack channel also occurs in response to pharmacological activation of the channel expressed in a mammalian cell line. Co-IP experiments were carried out using extracts of HEK cells expressing either the wild-type rat Slack or the *Slack-R455H* mutant channel, treated with or without the Slack channel activator niclosamide^25,32,33^. Transfection with Slack constructs did not alter the levels of FMRP, p-FMRP and CYFIP1 in cell lysates (**Fig. 2A,D**). Both in cells expressing the wild-type and *Slack-R455H* mutant channels, however, treatment with niclosamide resulted in higher levels of FMRP, p-FMRP and CYFIP1 bound to the Slack channel (**Fig. 2B,D**), but lower levels bound to eIF4E, (**Fig. 2C,D**). Collectively, our results indicate that pharmacological activation of Slack channels and the constitutively active Slack mutation trigger translocation of FMRP/CYFIP1 from eIF4E to the channel complex.

**Figure 2:**
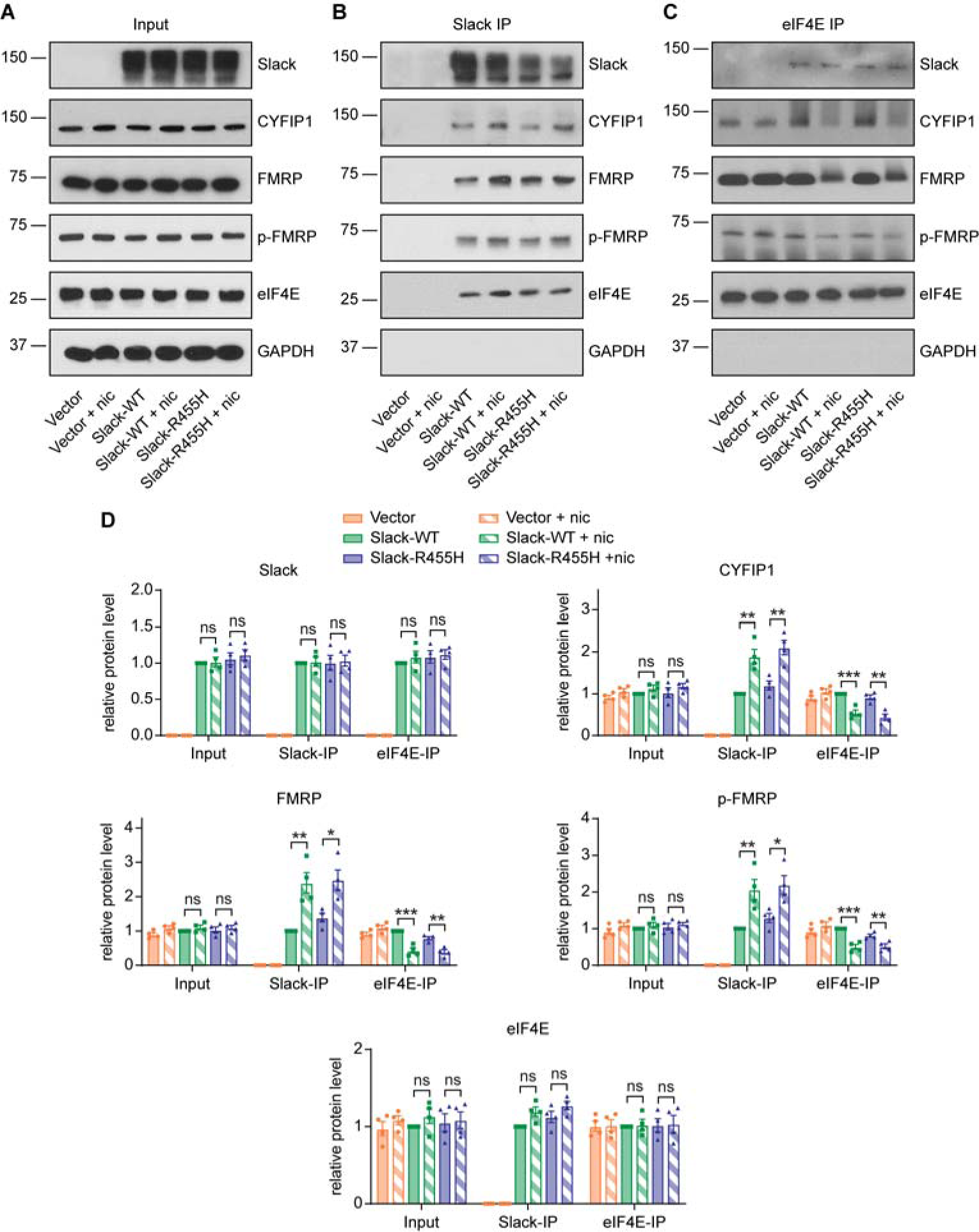
Activation of Slack channels triggers translocation of the FMRP/CYFIP1 complex. (A) Western blotting showing protein expression in HEK cells transfected with vector control, rat wild-type Slack (*Slack-WT*), or *Slack-R455H* treated with/without niclosamide (nic, 5 µM, 30 min). (B, C) Immunoprecipitation of Slack (B) or eIF4E (C) in HEK cells treated with/without niclosamide (5 µM, 30 min). (D) Quantification of western blots in a-C. Input, Slack IP, and eIF4E IP samples are normalized by GAPDH, Slack, and eIF4E levels, respectively, followed by normalization to *Slack-WT* level (n=4 replicates, two-way repeated measures ANOVA followed by multiple comparisons with paired t-test with Holm-Sidak correction). Data are expressed as mean ± SEM. *\*p*<0.05, *\*\*p*<0.005, *\*\*\*p*<0.0005

### Slack-R455H increases protein expression of a β-actin reporter construct

To examine the effect of translocation of the FMRP/CYFIP1 complex to Slack channels on translation, we used a reporter construct, dendra2-actin, which contains the coding region of the green fluorescent protein dendra2 flanked by the β-actin 5’ and 3’ UTR in a pCS2 vector (**Fig. 3A**). This reporter was selected because β-actin mRNA is a known binding target of FMRP^34^ and β-actin is rapidly translated in response to neuronal stimulation^35^. Dendra2 fluorescence can be irreversibly photoconverted to red fluorescence or photobleached using a 405 nm or 488 nm laser respectively^36,37^. This construct contains a non-functional myristylation tag (myr) that would normally target the protein to the cell membrane. This non-functional tag was removed for a subset of experiments described later.

**Figure 3:**
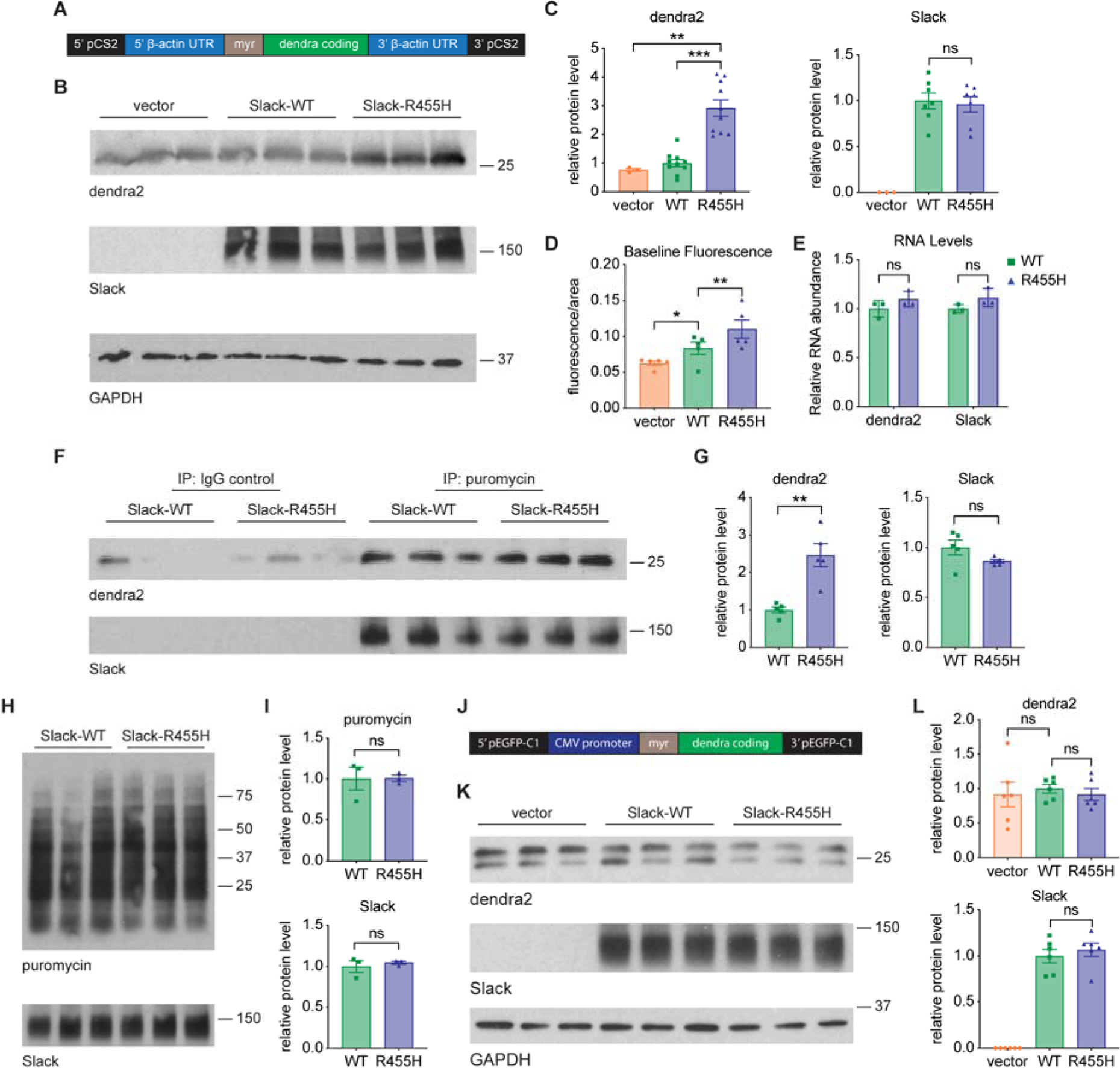
*Slack-R455H* mutation increases translation of **β**-actin reporter construct. (A) Schematic of β-actin translation reporter construct: fluorescent dendra2 protein flanked by 5’ and 3’ UTR of β-actin. Contains non-functional myristylation (myr) tag. (B) Western blotting showing protein expression in HEK cells 24 hours after co-transfection of dendra2-actin with vector control, rat wild-type Slack (*Slack-WT,* WT), or *Slack-R455H* (R455H). (C) Quantification of western blots in B (n=3-10 replicates). (D) Quantification of baseline fluorescence intensity of HEK cells 24 hours after co-transfection of dendra2-actin with vector control, *Slack-WT*, or *Slack-R455H* (n=5 plates, 6 FOV per plate, 3-22 cells per FOV). (E) qPCR of dendra2 and Slack mRNA levels 24 hours after co-transfection of dendra2-actin with *Slack-WT* or *Slack-R455H* (n=3 replicates). (F) Immunoprecipitation of puromycin in HEK cells following puromycin incubation 24 hours after co-transfection of dendra2-actin with *Slack-WT* or *Slack-R455H*. Shows actively translating protein. (G) Quantification of western blots in F (n=5 replicates). (H) Western blotting of HEK cells after puromycin incubation 24 hours after transfection with *Slack-WT* or *Slack-R455H*. Shows global translation levels. (I) Quantification of western blots in H (n=3 replicates). (J) Schematic of sub-cloned dendra2 reporter construct inserted into pEGFP-C1 vector without β-actin UTR. (K) Western blotting showing protein expression in HEK cells 24 hours after co-transfection of dendra2-C1 with vector control, *Slack-WT*, or *Slack-R455H*. (L) Quantification of western blots in K (n=6 replicates). (C, G, I, L) Western blotting without immunoprecipitation is normalized by GAPDH (C, L) or Slack (I) levels. All western blotting is then normalized to *Slack-WT* level (C, G, I, L). All comparison used unpaired t-test. Data are expressed as mean ± SEM. *\*p*<0.05, *\*\*p*<0.005, *\*\*\*p*<0.0005. See also Figure S1.

We first compared levels of dendra2-actin in HEK cells transiently co-expressed with either wild-type rat Slack or the gain-of-function mutant *Slack-R455H*. While wild-type Slack had no effect relative to an empty vector, *Slack-R455H* significantly increased levels of dendra2 protein detected on western blots (**Fig. 3B,C**). Slack protein levels themselves were unchanged between wild-type Slack and *Slack-R455H* expressing cells, ruling out preferential translation of dendra2 due to a poorly expressed *Slack-R455H* construct (**Fig. 3B,C**). We also measured baseline dendra2 fluorescence levels, as increased reporter expression should lead to increased fluorescence. Dendra2 baseline fluorescence was significantly increased when co-expressed with wild-type Slack compared to an empty vector and further increased when expressed with *Slack-R455H* (**Fig. 3D**). The discrepancy between the effects of wild-type Slack on dendra2-actin expression is likely due to altered sensitivity in measuring fluorescence levels compared to protein levels, as the fluorescence assay examines individual transfected cells, while the western blotting measures bulk protein including untransfected cells.

Co-expression with Slack could regulate dendra2-actin levels at several stages of protein synthesis and degradation. To rule out transcriptional regulation, quantitative real-time PCR (qPCR) was performed to examine mRNA levels for both dendra2-actin and Slack. No differences in mRNA levels were observed when dendra2-actin was co-transfected with wild-type Slack or *Slack-R455H*, confirming that transcription does not play a role in *Slack-R455H* regulation of dendra2-actin protein levels (**Fig. 3E**).

### Slack-R455H mutation specifically increases translation of β-actin reporter construct

We next examined whether *Slack-R455H* regulates the translation of dendra2-actin mRNA using a puromycin incorporation assay. Puromycin is a mild translation inhibitor that covalently binds to the nascent peptide strand at the ribosome, causing its release^38^. Incubating cells with puromycin prior to protein extraction allows the labeling of proteins that are being actively synthesized and can be used to measure global translation levels. We combined this technique with coimmunoprecipitation to detect the translating population of specific proteins. We found that levels of actively translating dendra2 are increased when expressed with *Slack-R455H* compared with wild-type Slack (**Fig. 3F,G**). Again, there were no significant changes in the rate of incorporation of puromycin into Slack.

Increased levels of dendra2-actin translation, with no change in levels of Slack, suggest that *Slack-R455H* specifically regulates dendra2-actin translation without affecting global translation. To directly confirm the specificity of regulation, we performed a traditional puromycin incorporation assay in the absence of dendra2-actin. The expression of *Slack-R455H* had no effect on global translation levels compared to wild-type Slack (**Fig. 3H,I**). While this result does not rule out additional targets of Slack translation regulation, it confirms that its regulation is limited in scope.

### β-actin UTR is the target of Slack-R455H regulation

The specificity of the effect of *Slack-R455H* in stimulating translation of dendra2-actin could, in theory, reside in sequences in either the β-actin UTR or the dendra2 coding region. To determine if regulation by *Slack-R455H* requires the β-actin UTR, we generated an alternate dendra2 construct (dendra2-C1), in which the coding region of dendra2 was inserted into an alternate vector (pEGFP-C1), which lacks the β-actin UTR (**Fig. 3J**). We found no difference in dendra2 protein levels when dendra2-C1 was co-expressed with an empty vector, wild-type Slack, or *Slack-R455H* (**Fig. 3K,L**), confirming that the β-actin UTR is a target of Slack regulation.

### Slack-R455H regulates translation initiation

A canonical mechanism by which the FMRP/CYFIP1 complex regulates translation of an mRNA is to control the rate of initiation by sequestering eIF4E^39^. Nevertheless, some proteins previously implicated in epilepsy and intellectual disability, including FMRP itself, can alter other steps in translation including ribosome stalling and translation termination^40-43^. These different mechanisms make different predictions about the amount of nascent protein product associated with ribosomes (**Fig. S1**). If *Slack-R455H* was to increase translation by relieving stalling during elongation or termination, levels of nascent dendra2-actin on ribosomes would decrease relative to those in cells expressing wild-type Slack or not expressing the channel (**Fig. S1A)**. In contrast, increased initiation of translation predicts increased levels of nascent dendra2-actin associated with ribosomes (**Fig. S1B**).

To differentiate between these two possibilities, we isolated ribosomes to measure the amount of bound dendra2-actin. Ribosomes were pelleted by centrifugation on a sucrose cushion according to a previously established protocol^44^. A validation experiment revealed that cytoplasmic proteins were exclusively found in the supernatant while ribosomal proteins were exclusively found in the pellet (**Fig. S1C,D**). Membrane proteins, however, also pelleted with the ribosomal fraction. While most dendra2 was in the cytosolic fraction, a small portion was found in the pelleted fraction, presumably ribosome-bound dendra2. However, because our original dendra2-actin construct contained a non-functional myristylation tag which normally targets protein to the membrane, we generated a construct without the myristylation tag (dendra2-Δmyr) to ensure that no fraction of dendra2-actin was targeted to membrane. When we performed ribosome isolation from cells co-expressing dendra2-Δmyr and Slack, we found that there was significantly more dendra2 bound to ribosomes in the presence of *Slack-R455H* than wild-type Slack (**Fig. S1E,F**). As in previous experiments, Slack levels did not change. These results are consistent with the model of **Fig. S1B**, in which Slack regulates the initiation of translation.

### Activation of wild-type Slack channels induces translation of **β**-actin reporter construct

To further test the effects of Slack activation on translation, we used the FRAP (Fluorescence Recovery After Photobleaching) technique to monitor dendra2-actin fluorescence in real-time. Dendra2 is irreversibly photoconverted to red fluorescence or irreversibly photobleached by 405 nm and 488 nm light, respectively (**Fig. 4A**)^36,37^. Because the rate of maturation of the dendra2 protein is fast, the rate of fluorescence recovery after the photobleaching of an entire cell provides a measure of the synthesis of new dendra2 protein (**Fig. 4B**). Although some previous studies with dendra2 have used photoconversion rather than photobleaching for this purpose^45,46^, we found in preliminary experiments that photobleaching caused less cytotoxic damage to the cells, while still allowing sufficient depletion of green fluorescence to observe recovery.

**Figure 4:**
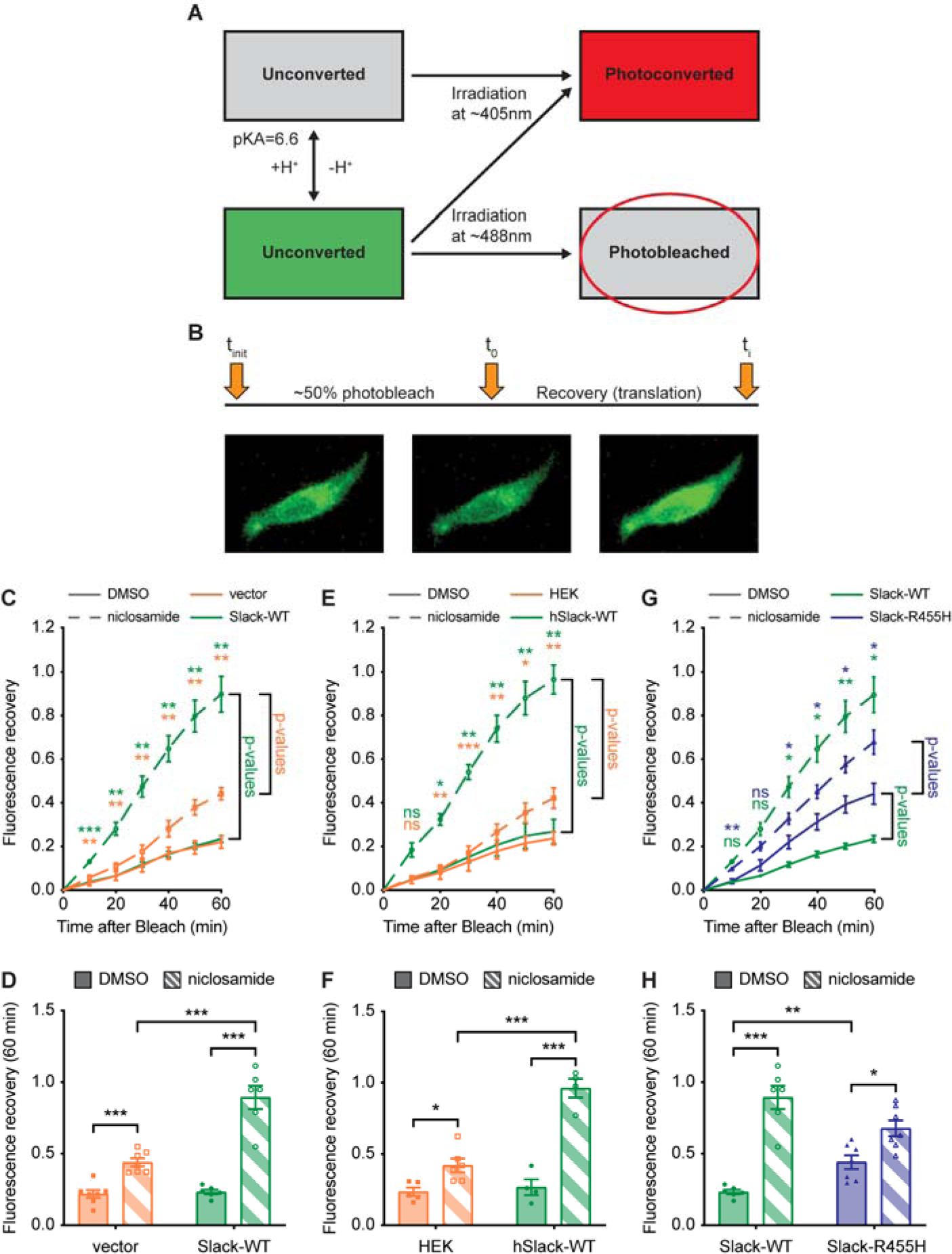
Slack activation increases translation of **β**-actin reporter construct. (A) Schematic of dendra2 fluorescence changes. (B) *Top:* Schematic of FRAP experiment protocol. *Bottom:* Example cell from FRAP experiments (*Left* = pre-photobleaching. *Center* = post-photobleaching. *Right* = post-recovery). (C-H) FRAP experiments with and without Slack activator niclosamide (10 µM, 30 min) in HEK cells. All cells were transfected with dendra2-actin 48 hours prior to imaging. (C, D) Vector control or rat wild-type Slack (*Slack-WT*) were co-transfected with dendra2-actin. (E, F) Only dendra2-actin was transfected into stable human Slack cells (*hSlack-WT*) or untransfected control cells. (G, H) *Slack-WT* or *Slack-R455H* were co-transfected with dendra2-actin. (C, E, G) FRAP time course. p-values represent differences between the groups indicated by the line right of the graph of the corresponding color (two-way repeated measures ANOVA followed by multiple comparisons with unpaired t-test). (D, F, H) Bar graph of 60-minute recovery time-point (unpaired t-test). All comparisons used Holm-Sidak correction (n=4-8 plates, 13-53 cells per plate). Data are expressed as mean ± SEM. *\*p*<0.05, *\*\*p*<0.005, *\*\*\*p*<0.0005. See also Figure S2.

We first co-transfected wild-type Slack with dendra2-actin into HEK cells. Co-transfection of the reporter with empty vector was used as a negative control. The time course of fluorescence recovery was determined using a two-photon microscope with automated software that obtains baseline images, identifies and photobleaches cells by 50%, and then obtains post-bleach images at 10-minute intervals. We compared the time courses of recovery in the presence or absence of the Slack channel activator niclosamide. In the absence of niclosamide, wild-type Slack had no effect on fluorescence recovery of dendra2-actin (**Fig. 4C,D**). In the presence of niclosamide, however, the rate of fluorescence recovery was greatly increased relative to negative controls. Niclosamide caused a slight increase in fluorescence in negative controls relative to DMSO, likely due to off-target effects, but these changes were much smaller than seen in the presence of wild-type Slack.

To determine whether the ability of Slack channel activation to stimulate synthesis of the reporter is conserved in human Slack channels, we repeated the above experiment using HEK cells stably expressing the wild-type human Slack channel (*hSlack-WT*). These cells were transiently transfected with dendra2-actin. HEK cells not expressing the channel were used as the negative control. As with the rat channel, *hSlack-WT* had no effect on dendra2-actin fluorescence recovery in the absence of niclosamide, but the rate of recovery was markedly stimulated by the channel activator (**Fig. 4E,F**). These data indicate that pharmacological activation of both rat and human Slack channels increases translation of the dendra2-actin reporter.

To determine whether K^+^ flux through the Slack channels is required for stimulation of dendra2-actin reporter synthesis, we used the non-specific channel pore-blocker quinidine ^47,48^, which inhibits Slack channel currents^33^. We carried out an additional FRAP experiment in HEK cells expressing *hSlack-WT* in the presence and absence of niclosamide and in the presence of both niclosamide and quinidine. We found that quinidine blocked the increase in dendra2-actin fluorescence recovery produced by the activator alone (**Fig. S2A,B**). While this suggests that K^+^ flux through the Slack channel pore may be required for activation of translation, quinidine is a general channel blocker with many potential off-target effects^47,48^, and definitive conclusions will require development of more selective pore blockers.

We next tested whether stimulation of dendra2-actin reporter synthesis is simply a consequence of increased K^+^ conductance. We compared the effects of Slack with those of the structurally related BK (K_Ca_1.1, KCNMA1, Slo1) channel. A CHO cell line that stably expresses the BK channel was transiently co-transfected with the dendra2-actin reporter. Treatment of these cells with the well-characterized BK channel activator NS1619^49^ had no significant effect on the rate of fluorescence recovery after photobleaching (**Fig. S2C,D**), suggesting that the increases in translation produced by Slack channels do not result simply from elevated K^+^ flux.

### Slack-R455H channels constitutively stimulate translation of **β**-actin reporter construct

We next carried out similar experiments using cells co-transfected with *Slack-R455H* and dendra2-actin. Consistent with the results of the biochemical experiments in **Figures 3** and **5**, the presence of the mutant channel significantly increased the rate of fluorescence recovery compared to wild-type Slack, even in the absence of a channel activator (**Fig. 4G,H**). Previous work, as well as that in **Figure 2**, has shown that some disease-causing mutant Slack channels can still be further activated by niclosamide^25^. Here, we found that treatment of cells expressing *Slack-R455H* with niclosamide did result in a further increase in fluorescence recovery. The degree of stimulation induced by niclosamide was, however, slightly reduced relative to wild-type Slack (**Fig. 4G,H**).

**Figure 5:**
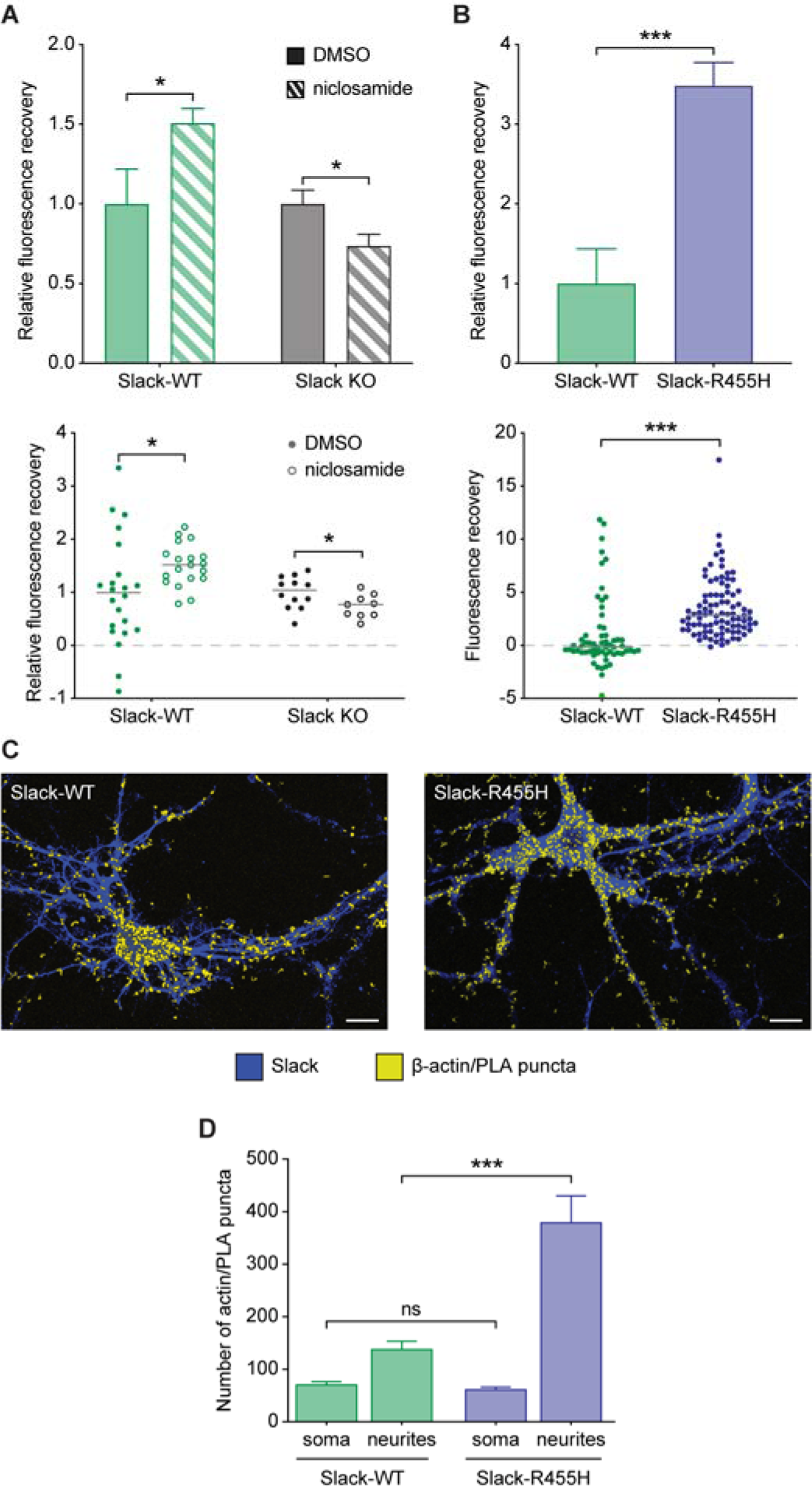
Endogenous neuronal Slack channels regulate **β**-actin translation. (A) Bar graph of FRAP experiment in primary mouse cortical neurons from *Slack-WT* mice or *Slack^-/-^* mice with and without Slack activator niclosamide (10 µM). Each data point (lower panel) represents an individual cell (n=9-22 cells). (B) Bar graph of FRAP experiment in primary mouse cortical neurons from *Slack-WT* or *Slack-R455H* mice. Each data point (lower panel) represents an individual cell (n=61-91 cells). (C) Representative PLA images of individual neurons from *Slack-WT* (*left*) or *Slack-R455H* (*right*) mice. (Blue = Slack, Yellow = β-actin/PLA puncta, Scale Bar: 10 μm). (D) Quantification of puncta number from PLA images in the soma or dendrites (n=17-33 cells). All comparison used unpaired t-test. Data are expressed as mean ± SEM. *\*p*<0.05, *\*\*p*<0.005, *\*\*\*p*<0.0005

### Endogenous neuronal Slack channels regulate **β**-actin translation

To determine if the regulation of translation by Slack channels that we characterized in HEK cells also occurs in neurons, we first carried out similar FRAP experiments using primary cortical neurons, which express Slack protein endogenously^4^. Control experiments were carried out using cortical neurons isolated from Slack knockout (*Slack^-/-^*) mice^27,50,51^. Neurons were transiently transfected with dendra2-actin as in the experiments with HEK cells, but bleaching and recovery were monitored using manual epifluorescence microscopy and with a single recovery time point. As in HEK cells, the addition of the Slack activator niclosamide significantly increased fluorescence recovery of dendra2-actin (**Fig. 5A**). In contrast, treatment of Slack^-/-^ neurons with niclosamide did not stimulate fluorescence recovery but produced a small decrease in recovery. The difference between the small off-target effects of niclosamide in HEK cells and cortical neurons may reflect differences in the regulation of β-actin translation in these cell types.

To determine if the constitutive stimulation of dendra2-actin translation by the *Slack-R455H* mutant that was detected in HEK cells also occurs in native neurons, we compared translation in wild-type neurons to that in neurons isolated from the heterozygous *Slack-R455H* mice described earlier. FRAP experiments were carried out on primary cortical neurons from both types of mice transiently transfected with dendra2-actin. As in HEK cells, the *Slack-R455H* mutation caused a significant increase in fluorescence recovery of dendra2-actin compared to that in wild-type neurons (**Fig. 5B**), demonstrating that the *Slack-R455H* phenotype of dysregulated translation occurs in cortical neurons.

Constitutive stimulation of the dendra2-actin reporter translation by *Slack-R455H* in neurons implies that the translation of endogenous β-actin should also be comparably increased. To test this directly, we carried out a proximity ligation assay (PLA) between puromycin and β-actin (**Fig. 5C,D**). This assay specifically identifies local β-actin synthesis by staining individual puncta of newly translated protein^52,53^. The number of puncta in a given region of a cell represents the number of β-actin transcripts that are being actively translated and serves as a measure of endogenous translation rate. These experiments were also carried out using cortical neurons from wild-type and *Slack-R455H* mice. We examined translation levels in the soma and neurites (within 100 μm of the soma) of cultured neurons independently because activity-dependent translation in neurons is often a highly localized process^54-58^. Indeed, while we found no difference in the number of PLA puncta in the soma of neurons from wild-type and *Slack-R455H* mice, neurons from *Slack-R455H* mice had 2-3-fold more puncta in neurites and therefore higher levels of β-actin translation (**Fig. 5C,D**). Our findings are in accord with other recent findings that K_Na_ channels are highly abundant in dendrites of cortical and hippocampal neurons^59^.

### FMRP and CYFIP1 are required for Slack activity-dependent translation

The recruitment of FMRP and CYFIP1 to the Slack channel complex, resulting in reduced levels of both proteins bound to eIF4E, provides a ready explanation for increases in translation upon channel activation. If this is the case, deletion of FMRP and CYFIP1 would unlink Slack from the translation machinery. To test this hypothesis, we suppressed FMRP and CYFIP1 expression in HEK cells using siRNA. Western blots revealed that the siRNAs are highly effective in reducing total levels of FMRP and CYFIP1, but do not affect levels of Slack protein (**Fig. 6A-D**).

**Figure 6:**
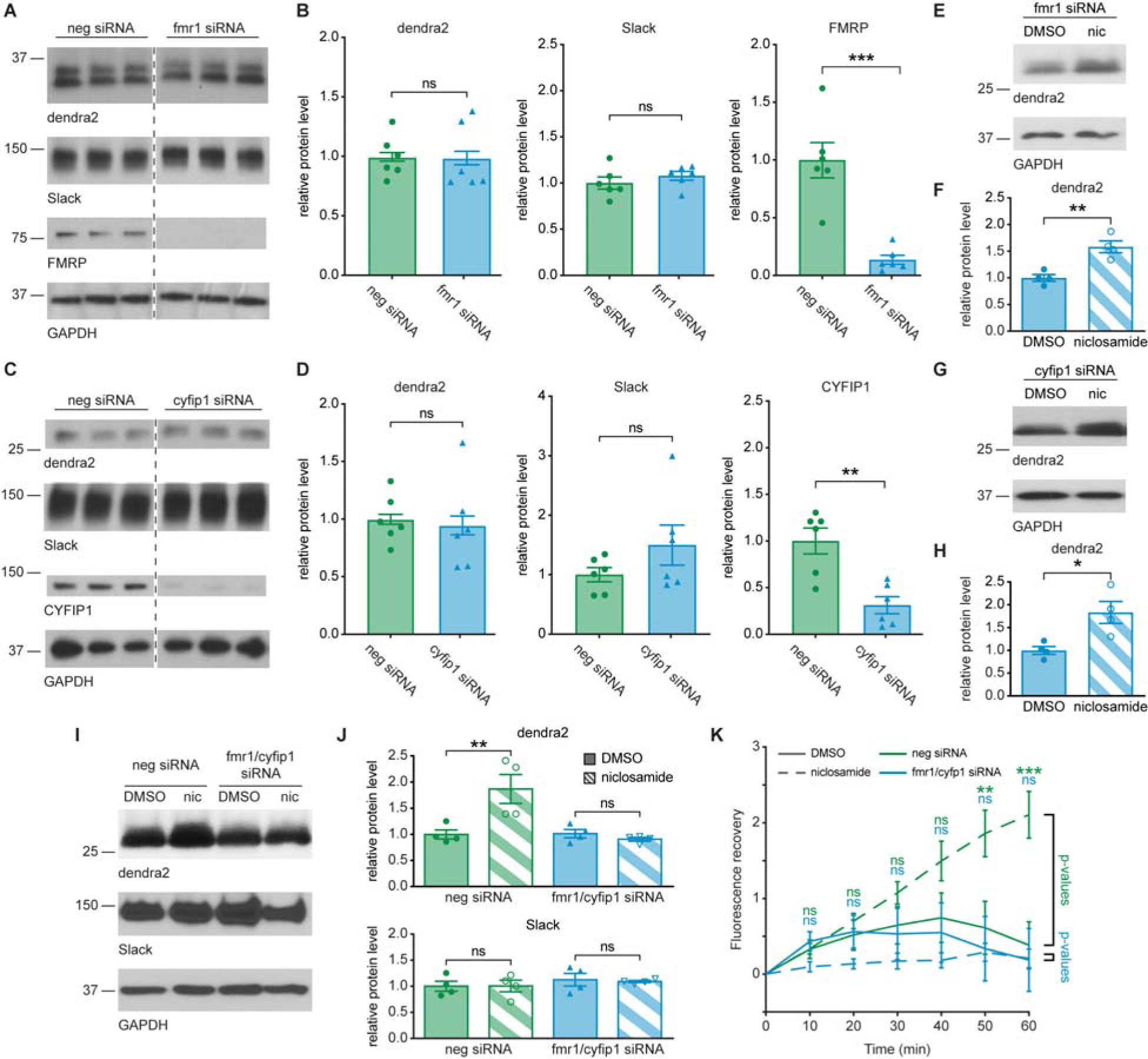
FMRP and CYFIP1 are required for Slack activity-dependent translation. (A, C) Western blotting demonstrating the effect of knockdown of FMRP (A) or CYFIP1 (C) on dendra2-actin expression levels in HEK cells. Cell extraction was carried out 24 hours after triple-transfection of dendra2-actin, *Slack-WT*, and *fmr1* siRNA cocktail (A) or *cyfip1* siRNA (C) or negative (neg) siRNA control. (B, D) Quantification of western blots in A and C, respectively. Protein levels were normalized to GAPDH level, followed by normalization to neg siRNA level (n=6 replicates, unpaired t-test) (E-J) Western blotting showing protein expression in hSlack-expressing HEK cells 48 hours after co-transfection of dendra2-actin with *fmr1* siRNA cocktail (E), *cyfip1* siRNA (G), or both *fmr1* and *cyfi*p1 (*fmr1/cyfip1*) siRNAs or negative siRNA control (I). Cells were incubated with or without Slack activator niclosamide (nic, 10 µM, 90 min) prior to protein extraction. (F, H, J) Quantification of western blots in E, G, and I, respectively (n=4 replicates). (F, H) Paired t-tests. (J) One-way ANOVA followed by multiple comparisons with paired t-test with Holm-Sidak correction. (K) Time course of FRAP experiment in hSlack-expressing HEK cells 48 hours after co-transfection of dendra2-actin with *fmr1/cyfip1* siRNAs or negative siRNA control, with and without Slack activator niclosamide (10 µM, 60 min). p-values represent differences between the groups indicated by the line right of the graph of the corresponding color (n=9-21 cells, two-way repeated measures ANOVA followed by multiple comparisons with unpaired t-test with Holm-Sidak correction). Data are expressed as mean *±* SEM. *\*p*<0.05, *\*\*p*<0.005, *\*\*\*p*<0.0005.

We co-transfected dendra2 with either *fmr1* or *cyfip1* siRNA, or with both siRNAs together into hSlack-expressing HEK cells. After 48 hours, cells were incubated with the Slack activator niclosamide or with DMSO alone for 90 min prior to protein extraction. Treatment with either *fmr1* or *cyfip1* siRNA alone did not prevent the increase in levels of dendra2 in response to the channel activator (**Fig. 6E-H**). Combined treatment with both siRNAs, however, eliminated the effect of niclosamide, while treatment with the negative control siRNAs had no effect on the ability of the activator to simulate dendra2 synthesis (**Fig. 6I,J**). Neither the siRNA for *fmr1* and *cyfip1* nor the negative control had any effect on levels of Slack. Finally, we also carried out FRAP experiments in hSlack-expressing HEK cells treated with *fmr1* and *cyfip1* siRNAs or the negative control, with and without niclosamide. Again, reducing both translation regulators fully eliminated the ability of niclosamide to stimulate the synthesis of the dendra2-actin reporter in the FRAP assay (**Fig. 6K**).

## DISCUSSION

We have found that activation of Slack Na^+^-activated K^+^ channels in cortical neurons, as well as in mammalian cell lines, results in the transfer of FMRP and CYFIP1 from the translation initiation factor eIF4E to the channel. The binding of the FMRP/CYFIP1 complex to eIF4E suppresses the initiation of translation^24^. Accordingly, we find that the increased binding of these two regulatory proteins to the channel is coupled to increases in the rate of translation initiation of a dendra2-actin reporter mRNA, and, in neurons, to increased synthesis of native β-actin itself. This effect can be recapitulated in cell lines expressing either rat or human Slack channels and is blocked by reducing expression of both FMRP and CYFIP1. Increased binding of FMRP/CYFIP1 to the channel, with higher levels of unbound eIF4E, are also produced by the gain-of-function mutation *Slack-R455H*, which results in constitutive activation of translation in both neurons and cell lines. Mice that are heterozygous for *Slack-R455H* have spontaneous seizures and persistent interictal spikes and the corresponding mutation in humans causes early onset epilepsy with severe intellectual disability^9,27^.

Slack channel activity is regulated by the binding of FMRP^10,11^. In neurons, intracellular injection of the peptide FMRP(1-298), rapidly increases the native Slack current and produces narrowing of action potentials^11^. In addition, single-channel open probability is increased when FMRP(1-298) is added directly to the cytoplasmic face of Slack channels in excised patches, further supporting a direct regulation of Slack by FMRP^10^. Our current findings suggest that the changes in neuronal firing pattern produced by these interactions are directly coupled to changes in translation of a subset of neuronal mRNAs. In addition, these findings raise the question of whether the interactions of FMRP with other ion channels^12-18^ influence protein synthesis or other biological processes regulated by FMRP. For example, although we did not find an increase in translation of dendra2-actin reporter with pharmacological stimulation of BK channels, changes in the availability of FMRP produced by binding to channels may influence its other functions such as localization of mRNAs or other aspects of cellular metabolism^43,60,61^.

Regulation of the translation of the dendra2-actin reporter by Slack requires the UTR regions of the β-actin gene. β-actin translation was selected as potential target of Slack regulation for several reasons. First, β-actin mRNA is a known binding target of FMRP^34^. As previous experiments demonstrated a functional interaction between Slack and FMRP, targets of FMRP binding are an obvious initial set of targets to examine^11^. β-actin is also known to be expressed both pre- and post-synaptically, where its translation is upregulated following synaptic stimulation^35,55^, as would be expected of targets of Slack activity-dependent translation^1,4^. Although we found that global translation levels are not affected by the *Slack-R455H* mutation, it is probable that other subsets of neuronal mRNA are also sensitive to Slack channel activation. For example, levels of the sodium channel Na_V_1.6 are substantially increased in the cerebral cortex of mice with the *Slack-R455H* mutation^62^. Further study will be required to determine whether Slack channel activation alters the translation of Na_V_1.6 and other mRNAs.

Our findings strongly suggest that severe intellectual disability, a hallmark of Slack-related epilepsies, does not result from the effects of the mutations on neuronal excitability alone. One of the conditions produced by gain-of-function mutations is Autosomal Dominant Nocturnal Frontal Lobe Epilepsy (ADNFLE). Patients with ADNFLE caused by Slack mutations have a very high chance of developmental delay and intellectual disability^7,63-65^. ADNFLE is, however, also caused by mutations in the nicotinic acetylcholine receptors CHRNA2, CHRNA4, or CHRNB2^64^. These patients almost always have normal cognitive development, despite identical seizure presentation to that caused by Slack mutations^64,65^. This strongly suggests that the disruption of Slack activity-dependent protein synthesis, rather than the seizures themselves, is a major factor that accounts for the severity of the effects of Slack mutations in humans. This conclusion is also consistent with the fact that dysregulation of translation is a recognized cause of intellectual disability^40-43,63,66^.

## ACKNOWLEDGEMENTS

This research was supported by NIH Grants NS102239 and DC 01919 to LKK and NIH Grant GM007324 to TJM.

## AUTHOR CONTRIBUTIONS

Conceptualization, T.J.M., J.W., and L.K.K.; Methodology, T.J.M, J.W., Y.Z., P.L., and R.C.; Software, T.J.M., S.N., and M.P.; Investigation, T.J.M, J.W., Y.Z., P.L., R.C., S.N., J.W., and M.P.; Resources, J.W.; Writing – Original Draft, T.J.M and L.K.K.; Writing – Review & Editing, T.J.M, J.W., Y.Z., and L.K.K; Funding Acquisition, T.J.M. and L.K.K.; Supervision, E.A.J and L.K.K.

## Supplementary Information

**Figure S1:**
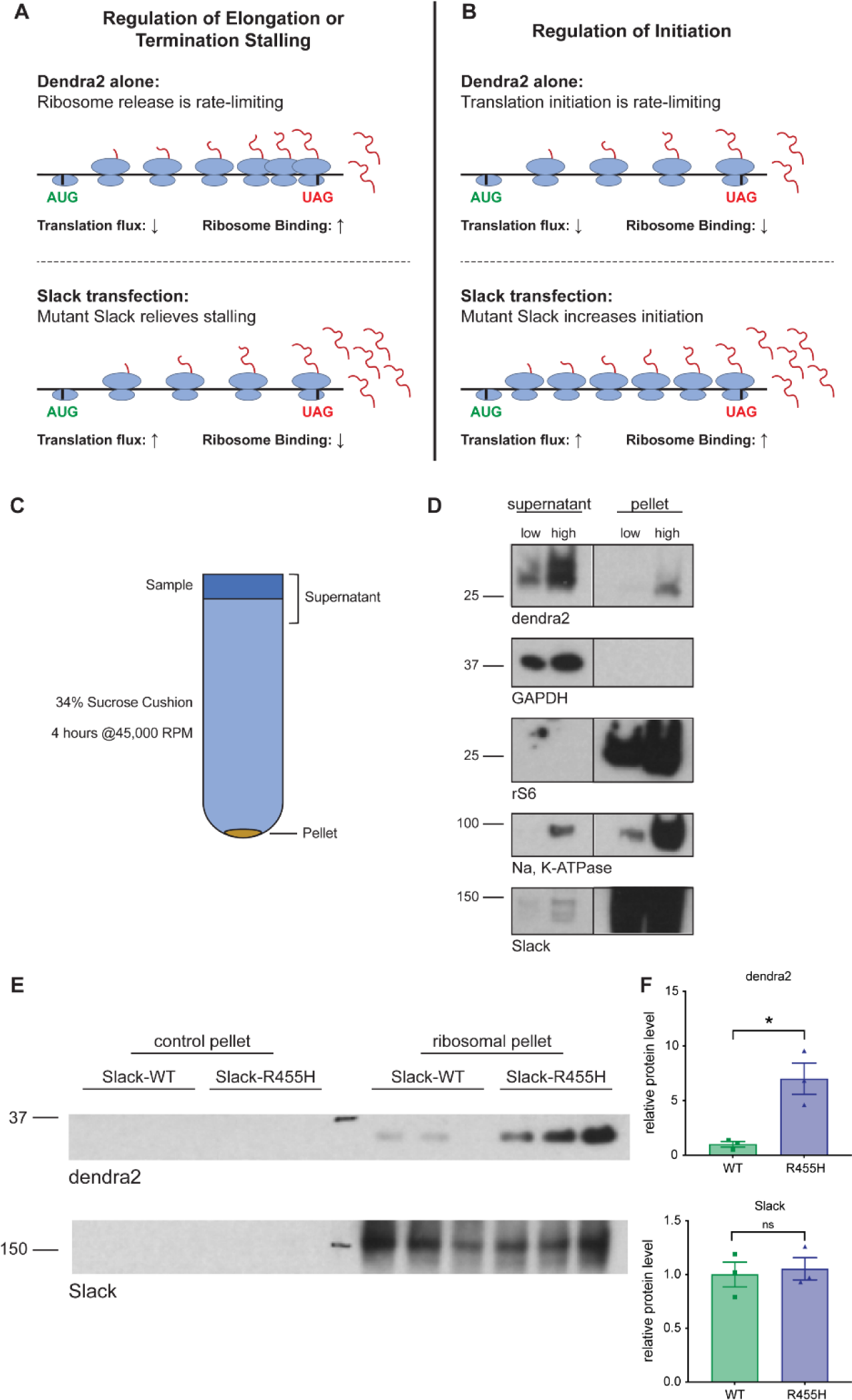
*Slack-R455H* regulates translation initiation, Related to Figure 3. (A) Model where Slack regulates translation through elongation or termination stalling. *Top*: In cells without Slack, ribosomes stall on dendra2 mRNA during elongation or termination. This leads to low total translation levels, but high ribosome binding. This requires a secondary mechanism that stalls ribosomes under baseline conditions, such as stalling through UPF3B or FMRP. *Bottom*: When mutant Slack is present, ribosome stalling is relieved, leading to increased total translation levels, but decreased ribosome binding. (B) Model where Slack regulates translation initiation. *Top*: In cells without Slack, translation occurs normally, but with low total translation levels and ribosome binding. This model does not require a secondary mechanism to explain the baseline condition. *Bottom*: Mutant Slack increases the rate of translation initiation. This leads to both increased ribosome binding and total translation levels. (C) Schematic of ribosome isolation protocol. 200 μL protein sample is loaded on top of 2 mL 34% sucrose cushion and centrifuged for 4.5 hours at 45,000 RPM. Validation samples were taken from the pellet and the supernatant for use in D. (D) Validation of ribosome isolation protocol. Pellet includes ribosomal proteins (rS6) and membrane proteins (Na, K-ATPase and Slack). Cytosolic proteins (GAPDH) are found only in the supernatant (top 400μL). Dendra2 is primarily found in supernatant, but a small fraction is found in the pellet. Paired lanes represent a 4x difference in input concentration of single sample. HEK cells were co-transfected with dendra2-actin and rat wild-type Slack (*Slack-WT*) 24 hours prior to protein extraction. (E) Western blotting showing ribosome binding of Δmyr-dendra2. HEK cells were co-transfected with Δmyr-dendra2 and *Slack-WT* or *Slack-R455H* 24 hours prior to protein extraction. Samples were run as described in C. Control pellet was obtained by re-centrifuging top 400μL of cushion to act as quality control. Dendra2 staining shows actively translating dendra2 still bound to ribosomes. Slack staining represents total Slack in cell membrane and serves as loading control. (F) Quantification of western blots in E. Dendra2 levels are normalized by Slack protein level followed by normalization to WT level. Slack levels are normalized to WT level (n=3, unpaired t test). Data are expressed as mean ± SEM. *p<0.05

**Figure S2:**
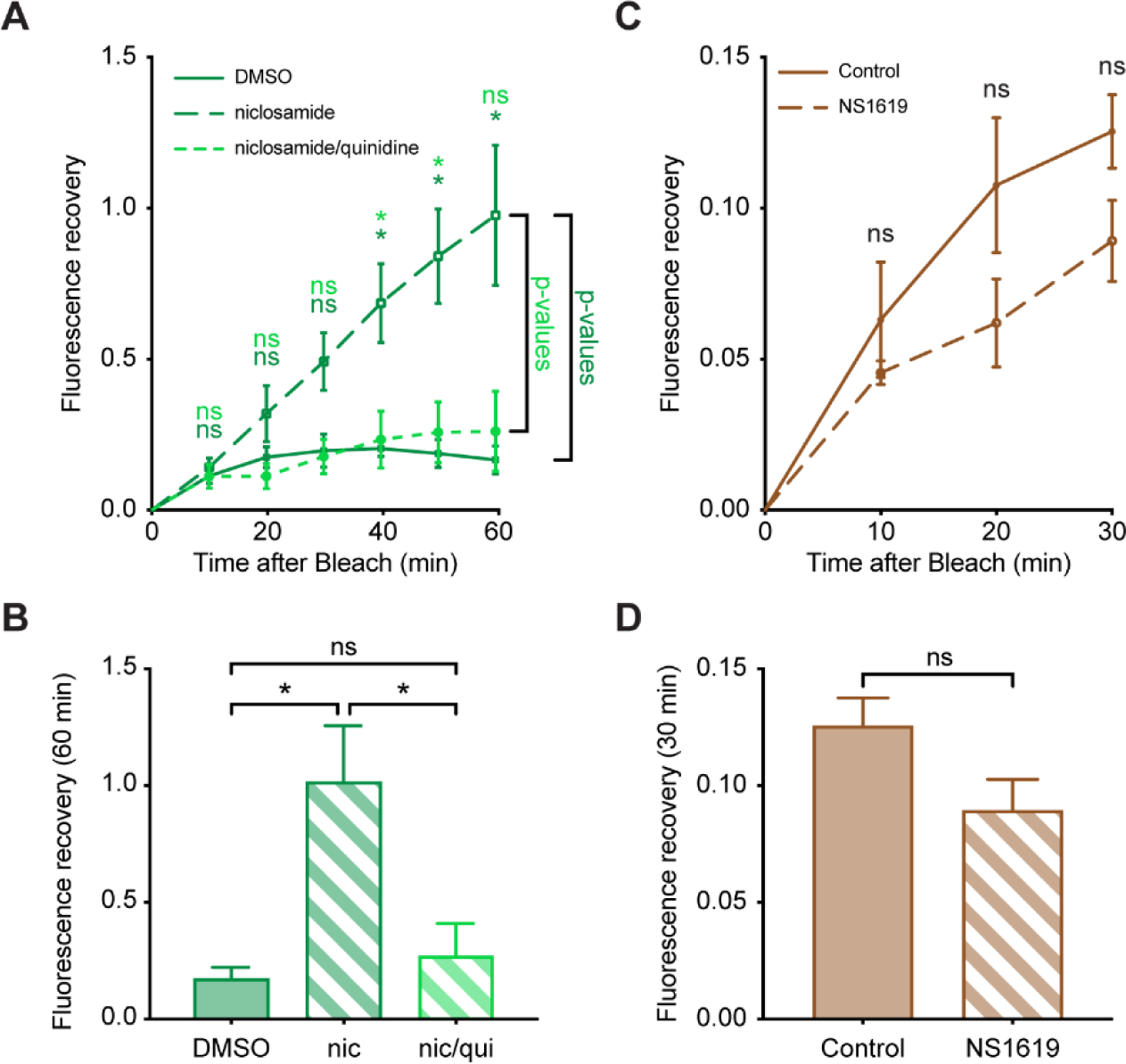
Slack regulation of translation requires ion conductance and is specific to Slack, **Related to** Figure 4. (A, B) FRAP experiment with and without Slack activator niclosamide (10µM, 30 min) and/or Slack pore-blocker quinidine (500µM, 30 min) in HEK cells stably expressing human Slack. All cells were transiently transfected with dendra2-actin 48 hours prior to imaging (n=4-6 plates, 1-12 cells per plate). (C, D) FRAP experiment with and without BK activator NS1619 (10µM, 30 min) in CHO cells stably expressing the BK channel. All cells were transiently transfected with dendra2-actin 48 hours prior to imaging (n=3-5 plates, 14-58 cells per plate). (A, C) FRAP timecourse. p-values represent differences between the groups indicated by the line right of the graph of the corresponding color (two-way repeated measures ANOVA followed by multiple comparisons with unpaired t-test) (B, D) Bar graph of 60-minute recovery time-point (unpaired t-test). All comparisons used Holm-Sidak correction. Data are expressed as mean ± SEM. *\*p*<0.05, *\*\*p*<0.005, *\*\*\*p*<0.0005

## RESOURCE AVAILABILITY

### Lead contact

Further information and requests for resources and reagents should be directed to and will be fulfilled by the Lead Contact, Leonard K. Kaczmarek.

### Materials availability

The *Slack^+/R455H^* mouse line is available from the Lead Contact, Leonard K. Kaczmarek, with a completed Materials Transfer Agreement.

### Data and code availability

- All data and the custom code used for semi-automated FRAP is available upon request.
- Any additional information required to reanalyze the data reported in this paper is available from the Lead Contact upon request.

## EXPERIMENTAL MODEL AND SUBJECT DETAILS

### Animals

Rodents were handled in accordance with protocols approved by the Yale University Institutional Animal Care Committee. Mice were housed on a 12/12 light/dark cycle with access to food and water ad libitum. Slack WT mice (*Slack-WT*) were C57BL/6J mice. Slack KO (*Slack^-/-^*) and *Slack-R455H* mice were generated as previously described(Lu et al., 2015; Quraishi et al., 2020). Genotyping was performed by ear clip biopsy-derived genomic DNA using established PCR protocols.

### Primary neuronal culture

Primary cortical neurons were prepared from P2-3 mice (for niclosamide experiments), from E16-17 mouse embryos (for knock-in mouse experiments), or 2-month old mice (immunoprecipitation experiments) as described previously with modifications specific for this study(Park et al., 2017). After isolation of cortex from neonatal or embryonic brains, neurons were dissociated and seeded (on coverslips inside a 6 well plate: 0.2E6 cells/6 well-plate) onto plates containing neurobasal medium supplemented with B27 (Gibco), 5% FBS (Gibco), L-glutamine, and antibiotics (Gibco). After 2 hours incubation, primary cultures were maintained in neurobasal medium without FBS in a 5% CO_2_ and 20% O_2_ incubator at 37°C. Subsequently, half the medium was replaced every 2 days. Neuronal cultures were transfected using Lipofectamine 2000 and experiments were performed at DIV14-21. Depending on the culture density 0.1 to 0.5 μg of dendra2 DNA vector was used for transfection.

## METHOD DETAILS

### Drugs

All pharmacological agents were obtained as dry compounds and dissolved in dimethyl sulfoxide (DMSO) (AmericanBio) to form 1000x stock solutions. Niclosamide (Santa Cruz) was diluted to a final concentration of 5 or 10 μM as noted, quinidine (Tocris Bioscience) was diluted to a final concentration of 500 μM, and NS1619 (MedChemExpress) was diluted to a final concentration of 30 μM.

### DNA constructs and subcloning

Rat-Slack-B-mCherry (*Slack-WT*), *Slack-R455H*, and mCherry-alone (empty vector) were engineered and characterized as described^6,67^. The pCS2-myrDendra2-Actb5’UTR3’ (dendra2-actin) construct was a gift from Dr. Deanna Benson (Icahn School of Medicine). myrDendra2-Actb5’UTR3’ subcloning into pEGFP-C1 vector (dendra2-C1) was performed with restriction enzyme cloning using NEBNext Ultra II Q5 Master Mix (New England BioLabs) using the following primers: Forward primer – ATA**ACTAGT**ATGGGCACGGTGCTGTC; Reverse primer – AAT**GGTACC**TAGAAGCATTTGCGTCGAGTCTT); bold represents restriction enzyme target sequences. Subcloning PCR product and pEGFP-C1 backbone were cut using Kpn1-HF and Spe1-HF restriction enzymes (New England BioLabs) and ligated using Quick Ligase Kit (New England BioLabs) according to manufacturer’s protocols. The myristylation tag of dendra2 was removed to generate pCS2-ΔmyrDendra2-Actb5’UTR3’ (Δmyr-dendra2) using the Q5 Site-Directed Mutagenesis Kit, substituting NEBNext Ultra II Q5 Master Mix for kit’s polymerase (Forward primer – AACACCCCGGGAATTAACC; Reverse primer – CATGGCGAACTGGTGGCG). All constructs were verified by DNA sequencing from Yale Keck facility.

### Cell culture and transfection

Human Embryonic Kidney 293T (HEK) cells (ATCC) were maintained in high glucose DMEM media (Gibco). Media was supplemented with 10% FBS (Millipore Sigma) and 1% Penicillin-Streptomycin solution (Millipore Sigma) incubated at 37°C with 5% CO_2_. HEK cells stably transfected with human Slack (*hSlack-WT*) were obtained as a gift from Autifony Therapeutics. *hSlack-WT* cells were cultured in media supplemented with 500 μg/mL G418 (Thermo Scientific). Cells were split every 3-4 days when confluent using TrypLE Express (Gibco). For all immunoblotting and imaging experiments, cells dishes were precoated with 0.002% poly-L-lysine (Millipore Sigma) for 15 minutes followed by 2x wash with water. Cells were plated on dish in 2 mL DMEM at a density such that they were confluent 48 hours after plating. For standard immunoblotting, immunoprecipitation, and imaging experiments, cells were plated in 35 mm dishes (HEK - 2.5E5 cells, *hSlack-WT* - 4E5 to 4.5E5 cells). For ribosome binding assay experiments, cells were plated in 60 mm dishes (2.5x all 35mm values; all further quantities listed are for 35 mm dishes unless specified). Transient transfection was performed 24 hours after plating using Lipofectamine 2000 (Thermo Scientific) according to manufacturer’s protocol. Cells were transfected with 500 ng dendra2 construct and/or 500 ng of Slack construct, as specified, in 500 μL optiMEM (Gibco) media with 5 μL lipofectamine.

Chinese Hamster Ovary (CHO) cells were maintained similar to HEK cells with the following differences: Culture media was IMDM (Invitrogen) supplemented with 10% FBS, 1% Penicillin-Streptomycin solution, and 1% HT supplement (Invitrogen). Cells were split approximately every 2 days. CHO cells were plated at 2.5E5 cells/dish in 35 mm dishes.

### Puromycin assay

For standard immunoblotting or immunoprecipitation experiments, puromycin incorporation protocols were identical. Prior to protein extraction, cells were incubated with 10 μg/mL puromycin, dihydrochloride (Millipore Sigma) for 10 minutes at 37°C. Cells were immediately placed on ice and protein extraction was begun.

### RNA interference

FMRP knockdown was performed using a silencing RNA (siRNA) cocktail made with equal parts of three Silencer Select siRNA against the *fmr1* gene (Thermo Scientific: s5315, s5316, s5317)^25^. CYFIP1 knockdown was performed using Silencer Select siRNA against the *cyfip1* gene (Thermo Scientific: s23242)^25^. Silencer Select Negative Control 1 (Thermo Scientific) was used as a control. For transfection, 50pmol siRNA of either cocktail or both together was co-transfected with DNA plasmids using Lipofectamine 2000 as described above. The negative controls used the same amount of total siRNA.

### Immunoblotting/Coimmunoprecipitation

Protein lysates from HEK cells or two-month-old mice frontal cortex were prepared using Pierce IP Lysis Buffer (Thermo Scientific) supplemented with cOmplete EDTA-free Protease Inhibitor Cocktail (Millipore Sigma) according to manufacturer’s protocol. Protein quantification was performed using Pierce BCA Protein Assay Kit (Thermo Scientific). For standard immunoblotting experiments or coimmunoprecipitation input controls, 7.5 μg protein was loaded onto precast 15-well 4-15% Mini-Protean TGX Stain Free Gels (Bio-Rad).

For coimmunoprecipitation experiments 187.5 μg protein was transferred to 1.5 mL Eppendorf tubes. Sample was precleared with 70 μL Pierce A/G agarose beads (Thermo Scientific) or Anti-IgY PrecipHen beads (AvesLabs) (50% v/v Pierce IP lysis buffer). Tubes were incubated at 4°C for 1 hour on rotator. Samples were centrifuged at 3,000 rpm at 4°C for 2 minutes. 75 μg of sample was incubated at 4°C overnight on rotator with 5 μg mouse anti-puromycin antibody (Millipore Sigma: MABE343), or rabbit anti-eIF4E (Abcam, ab33766), or chicken anti-Slack IgY antibody, or IgG (Santa Cruz: sc-2025) or IgY (AvesLabs) control antibodies. 30 μL beads were added to samples, which were then incubated at 4°C for 2 hours on rotator. Samples were centrifuged at 3,000 rpm at 4°C for 1 minute to collect beads, which were washed 7 times with Pierce IP Lysis Buffer. Protein was eluted by incubation with 50 μL 2x Laemelli Buffer (Bio-Rad) at room temperature for 30 minutes on rotator. Mini-protean gel was loaded with 15 μL of eluted sample.

For all immunoblotting experiments, gels were processed and blots were developed as previously described^11^. Primary and secondary antibodies were used at the following concentrations: anti-dendra2 1:5000 (OriGene: TA150090), anti-Slack 1:5000 (Aves labs: custom), anti-puromycin 1:1000 (Millipore Sigma: MABE343), anti-FMRP 1:2000 (Millipore Sigma: MAB2160), anti-p-FMRP 1:1000 (Thermo Scientific: PA5-35389), anti-CYFIP1 1:500 (Santa Cruz: sc-49932), anti-eIF4E 1:1000 (Abcam, ab33766), anti-GAPDH 1:1000 (Santa Cruz: sc-32233), anti-rabbit-HRP 1:1000 (Thermo Scientific: 32460), anti-mouse-HRP 1:1000 (Thermo Scientific: 32430), anti-goat 1:10,000 (Thermo Scientific: 31402), anti-chicken-HRP 1:20,000 (Jackson Laboratories: 703-035-155). Wash steps were performed with three 10-minute washes. When multiple antibodies were stained successively on the same blot, blots were stripped as necessary with 5mL Restore Plus Western Blot Stripping Buffer (Thermo Scientific) followed by three 5-minute washes. Protein bands were analyzed with ImageJ (NIH) software. Where applicable, blots were normalized to GAPDH or Slack levels followed by normalization to the level of the denoted control group, as described in the figure legends. Both normalization steps were performed on a per-blot basis. When individual blots contained one sample of each condition (**Fig. 1, 2, 6E-J**), paired t-tests were performed. When individual blots contained multiple samples of each condition (**Fig. 3, 6A-D, Fig. S1**), unpaired t-tests were performed.

### Imaging experiments

For HEK and CHO cell imaging experiments, cells were replated 12 (baseline fluorescence) or 36 (FRAP) hours after transfection. Cells were rinsed twice with phosphate buffered saline (PBS) followed by a 5s rinse with 0.5mL TrypLE Express. Cells were replated onto fresh 35 mm dishes coated with poly-L-lysine (as described above) at a density of 1.5E5 cells/dish in 2 mL high-glucose DMEM. Cells were imaged 12-24 hours after replating. Media was changed to 2 mL phenol red-free DMEM media (Gibco) supplemented with 25 mM HEPES and 4 mM L-glutamine with any denoted drugs 30 minutes (except where noted) prior to imaging.

HEK and CHO cell imaging experiments were performed with semi-automated imaging using a Scientifica Hyperscope multi-photon system using custom-built Matlab scripts. For baseline imaging experiments the only active user input is initial focal plane selection. The program selects six arbitrary fields-of-view (FOV). For FRAP experiments, the user manually selects FOV. One to two FOV are imaged per plate, so manual selection is desired to ensure FOV contains enough cells without any over-bright cells or debris. Once FOV are selected, program takes initial Z-stack image of FOV and automatically segments image to identify all cells from sum-intensity projection of Z-stack image. In FRAP experiments, program photobleaches identified cells by about 50%, and takes a post-bleach Z-stack image. Bleaching is performed using a high laser power and high-density imaging. These steps are performed successively for each FOV. After the initial imaging and bleaching, microscope returns to each FOV at set time intervals taking a time course of recovery Z-stacks. For all automated imaging, fluorescence levels are automatically calculated for all cells and FOV. Range for Z-stack is determined by identifying focal plane edges past which total FOV fluorescence is below a threshold value. This automation allows significant imaging throughput, prevents user-generated bias in both focal plane selection and cell detection, and maximizes reproducibility. Imaging parameters are as follows: acquisition system – GalvoImaging; laser wavelength – 960nm; beam power – 20%, PMT gain – 600, image resolution – 1024 pixels; frame/slice – 4; slice number – 5; focus threshold – 600 A.U.; bleach power – 40%. Cells were filtered to include only healthy bleached cells. HEK cell inclusion criteria were as follows: min bleach: 20% baseline reduction; max bleach: 75% baseline reduction; min fluorescence (at t=0): 25 A.U.; max fluorescence (at t=0): 2000 A.U.; min recovery (at t=60): 0 A.U.). Quinidine and siRNA FRAP experiment used the following modifications due to laser recalibration and software updates. Quinidine: beam power – 25%; bleach power – 85%; manual segmentation (for bleaching only). siRNA: beam power – 25%; No bleaching (does not alter observed recovery, as seen with negative control siRNA), max timepoint 1 decrease – 50% baseline reduction (removes cells with large fluctuations), recover time point extremes – -75%, 400% (removes outlier cells). Due to differences in CHO cell bleaching and fluorescence levels, CHO cell inclusion criteria were different as follows: min bleach: not used; min fluorescence (at t=0): 20 A.U.; min recovery (at t=30): 0 A.U.

FRAP experiments in neurons were performed manually using a Zeiss Axiovert 200 epifluorescence microscope 24 hours post-transfection. Drug was directly added to media prior to imaging. After taking baseline image, cells were bleached for two minutes at full laser power, after which post-bleach image was taken. Recovery period for neurons was 5 minutes. Fluorescence intensity (center of the soma) was analyzed using ImageJ software. No inclusion criteria were used for neuron FRAP as recovery was calculated for individual cells at a single time point and fit a normal distribution (**Fig. 5A,B, lower panels**).

### Quantitative real-time PCR

For qPCR experiments, mRNA was isolated using RNeasy Mini Kit (Qiagen) and cDNA was generated using the ProtoScript II Reverse Transcription Kit (New England BioLabs). qPCR was performed on a LightCycler 480 real time PCR system (Roche) using PowerUp SYBR Green Master Mix (Thermo Scientific) according to manufacturer’s instructions. RNA levels for each condition were calculated using the single-delta CT method with GAPDH as the normalizing gene. Data for dendra2 and Slack represents the mean mRNA level of two independent primer pairs for a given sample.

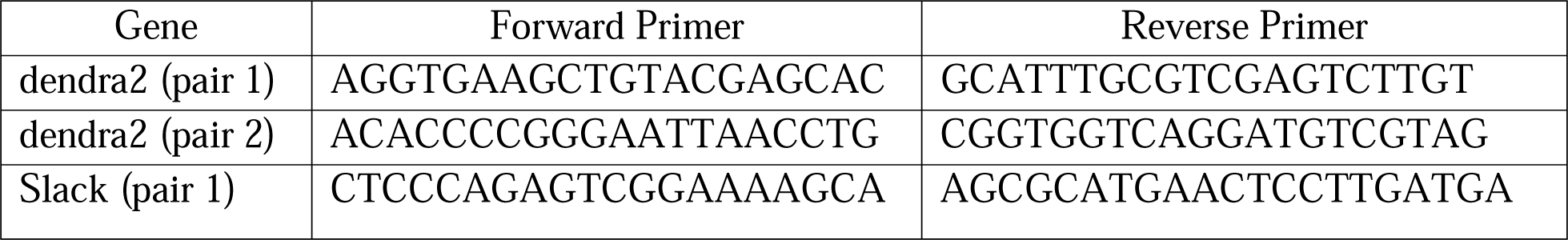
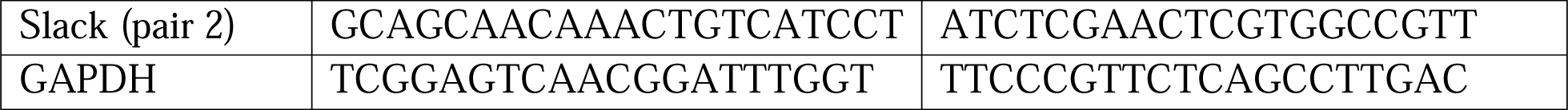

### Ribosome Binding Assay

Ribosome binding assay is adapted from previously published ribosome profiling protocol^44^. Above protein extraction protocol was used with the following changes: Cells were washed 3x with 2 mL ice cold PBS spiked with 100 μg/mL cycloheximide (CHX) (Sigma). Lysis buffer was replaced with 500 μL polysome lysis buffer (in mM except as noted): 20 Tris pH7.4, 150 NaCl, 5 MgCl_2_, 1 DTT, 100μg/mL CHX, 1x Protease Inhibitor Cocktail, 1% Triton X-100. Incubation at 4°C was not performed on rotator. Cells were triturated cells ten times through a 26-gauge needle after 4°C incubation. Clarifying centrifugation was changed to a 10 minute 5,000G centrifugation followed by a 20,000G centrifugation. Finally, samples were snap frozen in ethanol and dry ice and stored at -80°C until use.

For polysome isolation step, samples were diluted to 420μg protein in 240μL polysome lysis buffer. 2μL of RNase 1_f_ (New England Biolabs) was added to samples and incubated at room temperature for 45 minutes under gentle agitation. Meanwhile, a 1M (34% w/v) sucrose cushion was prepared (in mM except as noted): 20 Tris pH7.4, 150 NaCl, 5 MgCl_2_, 1 DTT, 1 sucrose, 100μg/mL CHX, 20U/mL SUPERase-In RNase Inhibitor (Thermo Scientific). 2mL of sucrose cushion was added to 13x51mm Ultra Clear tubes (Beckman). After incubation, 200μL of sample was gently added on top of sucrose cushion. Additional polysome lysis buffer was added to tubes to equalize mass (± 5mg). Ribosomes were pelleted by centrifugation in an SW 55 Ti rotor for 4.5 hours at 45,000 rpm. These conditions result in equal sedimentation to the conditions used in McGlincy et. al.^44^

After centrifugation, cushion fractions were saved as described in results. The pellet was extracted by adding 50μL 2x Laemelli Buffer and incubating for 10 minutes at 37°C. Tubes were briefly vortexed and pellet fraction was transferred to a microcentrifuge tube. Centrifugation process was repeated with another 2mL sucrose cushion and 200μL of top reserved fraction in fresh tubes, which served as a specificity and contamination control. Procedure for preparing input control samples and performing immunoblotting are as described above.

### Newly synthesized Actin-PLA labeling

Detection of newly synthesized proteins by proximity ligation assay was carried out using anti-puromycin antibody in combination with protein-specific antibodies and detection using Duolink reagents (Sigma) according to the manufacturer’s recommendations. Briefly, disassociated cortical neurons (DIV11) were treated with 10 μg/mL puromycin for 20 min and fixed for 20 min. After 3 times washing with PBS, cells were permeabilized for 15 min with 0.5% Triton X-100 in PBS, blocked in 4% goat serum in PBS for 1 h at 37°C. Cells were co-incubated with mouse anti-Puromycin antibody (Kerafast Equation 0001, 1:100) and rabbit anti-actin antibody (Cat# A2066, Sigma, 1:100) at 4°C overnight. PLA probes mouse PLA^plus^ and rabbit PLA^minus^ were used for the puro-actin PLA procedure as described previously^52,53^ and a far-red fluorescence-labeled oligo (Duolink Detection Reagents FarRed) was used for detecting the newly synthesized actin protein.

After PLA labeling, cells were post-fixed in 4% PFA at room temperature for 10 min, and washed with PBS for detection of the Slack expression immunostaining with the primary antibody anti-Slack (1:500, Aves labs: custom) and secondary antibody Alexa-Fluor 488 anti-chicken (1:500, Thermo Scientific: A-11039) as described previously^70^. Images were acquired using a Leica SP5 confocal microscope with a 100x magnification. Z-series stack confocal images were taken. All confocal images were processed with ImageJ. Actin/PLA puncta were counted on the soma and on neurites within 100 μm of the soma. If there were two or more closely spaced somata for which it was not possible to attribute specific neurites to a given soma, the somatic puncta were counted separately and all of the associated neuritic puncta were counted and divided equally among the somata.

## QUANTIFICATION AND STATISTICAL ANALYSIS

Data analysis, imaging processing, and graphing were performed using Microsoft Excel (Microsoft), MATLAB (MathWorks), and Prism (GraphPad). All data are expressed as mean ± SEM. Where noted, significance between groups was first calculated by two-way repeated measure ANOVA. All individual statistical comparisons were calculated by two-tailed Student’s t-test (paired or unpaired as applicable) with correction for multiple comparisons using the Holm-Sidak method when applicable.

